# Demographic history shapes forest tree vulnerability to climate change

**DOI:** 10.64898/2026.03.10.710859

**Authors:** Thomas Francisco, Isabelle Lesur-Kupin, Carlos Guadaño-Peyrot, Sanna Olsson, Marija Kravanja, Marjana Westergren, Sara Pinosio, Thibaut Capblancq, Giovanni G. Vendramin, Katharina B. Budde, Lene R. Nielsen, James M. Doonan, Delphine Grivet, Elia Vajana, Juliette Archambeau, Andrea Piotti, Santiago C. González-Martínez

## Abstract

Demographic history is expected to play a central role in shaping population vulnerability to climate change through its lasting effects on effective population sizes and genetic connectivity. However, existing studies report contrasting outcomes, and the consequences of alternative demographic histories have seldom been assessed concurrently across multiple taxa. Here, we analysed population genomic data from six of the major European forest tree species to address this gap. Across species, genetic isolation reduced adaptive potential, as reflected in lower standing genetic diversity. Greater genetic differentiation was also associated with increased maladaptation to climate, whereas the purge or accumulation of deleterious mutations depended on the severity of past demographic events. Finally, the beneficial effects of gene flow were evidenced across the six species by more optimal climate adaptation in highly connected populations. Altogether, our results provide valuable insights into how genetic differentiation, reflecting the combined effects of genetic drift and limited historical gene flow, influences current vulnerability of forest tree populations to climate change.

## Introduction

Trees are foundation species in many terrestrial ecosystems, supporting remarkable biodiversity, regulating the climate and providing other vital ecosystem services on a global scale^1^. Due to their long generation time and sessile nature, trees are particularly exposed to environmental hazards and gradual changes. An increasing body of literature indicates that they will not be able to adapt and/or disperse at rates matching projected climate change^2–4^ and forests worldwide are already displaying signs of stress and decline^5,6^. The way in which populations respond to environmental changes depends on the intensity of the stress they experience, their sensitivity to this stress and their capacity to adapt to the new conditions, which together determine their vulnerability to a changing environment^7,8^.

Genetic metrics are increasingly being used to assess vulnerability to climate change, as they capture impacts and responses at the intraspecific level^9–11^. Genetic diversity originates diverse phenotypes, some of which may confer advantages in shifting climate, thus forming the basis of the evolutionary potential of populations and, possibly, different patterns of local adaptation^12^. Although local adaptation is common in plants^13^, the degree of local adaptation can be population-specific^14^. Both adaptation lags to current climate and pre-adaptation to future climate can influence the extent to which a population will be affected by climate change^15,16^. Recent advances in ecological genomics have introduced new metrics to quantify the extent of climate adaptation in populations (e.g., the genomic discrepancy index, GDI^17^). Genetic variation between populations resulting from different rates of accumulation and purging of deleterious mutations, can also shape their sensitivity to climate change. Indeed, levels of genetic load, defined as the reduction of fitness in a population due to deleterious mutations, can further constrain a population’s ability to persist under shifting climatic pressures^18^. Additionally, inbreeding depression has been reported to be particularly high in gymnosperms compared to angiosperms^19,20^, as evidenced, for example, by the high number of lethal equivalents affecting tree fitness observed in conifers^20–24^. Overall, regardless of their impact on current fitness or adaptive potential, population attributes such as genetic diversity, climate adaptation, or genetic load are shaped by demographic processes, and are therefore neither static through time nor uniform across populations^23–27^.

Over evolutionary timescales, tree populations have been profoundly shaped by repeated glacial cycles, large-scale environmental shifts and, more recently, land-use changes. These events triggered large distributional shifts that came with important variation in population size and connectivity, and thereby shaped the genetic makeup of populations. The impact of demographic history on wild populations is widely recognised, primarily through its effects on effective population sizes, gene flow and range limits^26,28,29^. However, its overall impact on tree species’ genetic diversity remains unclear and requires broader investigation across species with contrasting demographic histories^30^. Demographic history can also affect a population’s potential for climate adaptation by influencing the amount of standing genetic variation available for natural selection and the patterns of adaptive or maladaptive gene flow^31–33^. Furthermore, genetic load is also impacted by demographic historical events, through the purging or accumulation of deleterious mutations^24,25^. Although well-established theoretical predictions on the link between demographic history and genetic load exist^34,35^, empirical studies have reported contrasting results^36–39^, highlighting the need for further research, particularly in forest trees.

In this study, we investigated how historical demographic processes may have shaped key components of population vulnerability, namely sensitivity and adaptive potential, in six ecologically and economically important tree species: four conifers (*Taxus baccata* L., *Pinus pinea* L., *Pinus pinaster* Ait. and *Pinus sylvestris* L.) and two angiosperms (*Fraxinus excelsior* L. and *Fagus sylvatica* L.). By integrating genomic data from 6,435 trees across 326 populations spanning most of the species’ distributions in Europe and surrounding regions, we estimated genetic diversity, genetic load, and current climate adaptation, and related these metrics to population-level genetic differentiation, used here as a proxy for demographic history. Finally, we interpreted these associations in the light of how historical demographic processes, including genetic drift and population isolation, may have influenced present-day sensitivity and adaptive potential to ongoing climate change.

## Results

### Population genetic differentiation reflects the species’ demographic history

The six tree species included in this study were chosen to follow a range of typical population sizes and levels of connectivity, likely resulting from the interplay between life-history traits and contrasting demographic histories. Population-level genetic differentiation (*F*_ST_) reflected well these patterns (Fig. 1). Populations of *T. baccata* and *P. pinea* consistently showed intermediate to high *F*_ST_ (median values of 0.151 and 0.380, respectively; higher than any other species), which may have arisen from their known reduced effective population size and connectivity, resulting from recurrent historical bottlenecks and long-term habitat contraction^40–42^. In contrast, *P. pinaster*, displayed greater variability in *F*_ST_ across populations, reflecting the presence of continuous and well-connected populations in parts of its range, as well as smaller and fragmented ones in other parts, likely resulting from contractions during glacial periods followed by post-glacial range expansions and reduced gene flow across gene pools^30,43^. Finally, *F. excelsior*, *F. sylvatica* and *P. sylvestris* populations displayed very low *F*_ST_, with only a few populations showing higher genetic differentiation (e.g., *F*_ST_ > 0.2 only in two *F. excelsior* and one *P. sylvestris* populations). Most populations of these species are known to be well-connected due to a combination of factors such as multiple glacial refugia, extensive post-glacial expansion and admixture, notable cold tolerance and efficient long-distance gene flow^19,30,43–51^. Nevertheless, smaller, fragmented populations can sporadically be found in few, mainly southern, parts of the range of these species due to environmental constraints or geographical barriers (*F. excelsior*^48^; *F. sylvatica*^52^; reviewed in Pyhäjärvi *et al*.^19^ for *P. sylvestris*).

**Figure 1.**
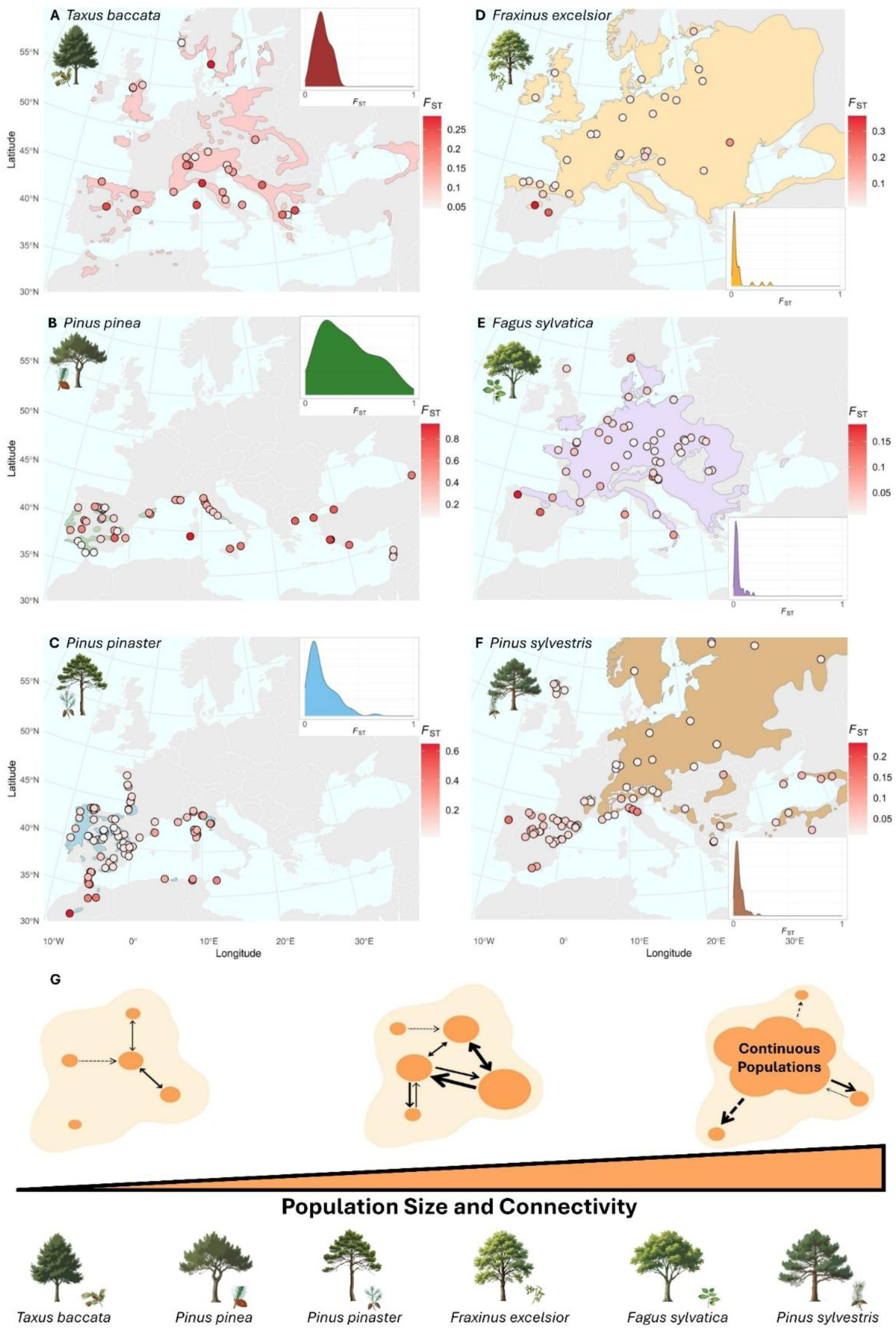
Description of the six tree species. **A-F** Geographical range and population-level genetic differentiation (*F*_ST_). Each circle represents a population, with colour intensity indicating the magnitude of *F*_ST_. Inserts show the density distribution of population-level *F*_ST_ values, represented on a common scale to facilitate visual comparison across species. Shaded areas depict the approximate species range^101^ for the geographic extent of latitude 30°N-60°N and longitude 10°W-37°E. Sampled populations of *P. sylvestris* occurring outside this range are not shown but can be found in Fig. S1. Notice that species such as *T. baccata*, despite having a large geographical range, have a scattered distribution while others (e.g., *P. sylvestris*) constitute large, stand-forming populations. **G** Conceptual illustration of the gradient in population size and connectivity represented by the six species under study^30,40–43,46,50^, which spans from species with predominantly small and isolated populations to species with large and well-connected populations. The schematic is inspired by^43^.

In addition, the geographical distribution of population-level *F*_ST_ varied among species. Highly differentiated populations of *T. baccata* and *P. pinea* showed no clear spatial pattern (Fig. 1A, B), with high *F*_ST_ even in populations located within the species’ core range. In contrast, the other species exhibited higher *F*_ST_ towards the range edges compared to their core populations (Fig. 1C-F; see also Theraroz *et al*. ^39^ and Guadaño-Peyrot *et al*.^53^).

### Key intraspecific components of climate change vulnerability vary across populations and species

Genetic diversity (*H*s) varied considerably within species, with distinct patterns emerging across species (Fig. 2A; but notice that direct comparisons of genetic diversity among species are not appropriate due to the use of different genotyping platforms; see Methods). At the within-species level, *P. pinea* exhibited the greatest among-population variability in *H*s, as evidenced by its relatively flat cumulative curve compared with the other species. The other two pine species also had higher among-population variability in *H*s than *T. baccata* and the two broadleaves, for which the cumulative curves were steeper. *H*s changed significantly with longitude in most species, increasing eastward in *T. baccata*, *F. sylvatica* and *P. sylvestris*, decreasing in *P. pinea* and *P. pinaster* and showing no significant trend in *F. excelsior* (Fig. S2A; Table S1). Additionally, *H*s increased significantly with latitude in *P. pinaster* and *P. sylvestris*, with southern range-edge populations of both species showing lower genetic diversity (Fig. S2B). These patterns likely reflect known glacial and post-glacial forest dynamics as reconstructed from genetic data and pollen records (e.g.,^39,42,44,54–58^).

**Figure 2.**
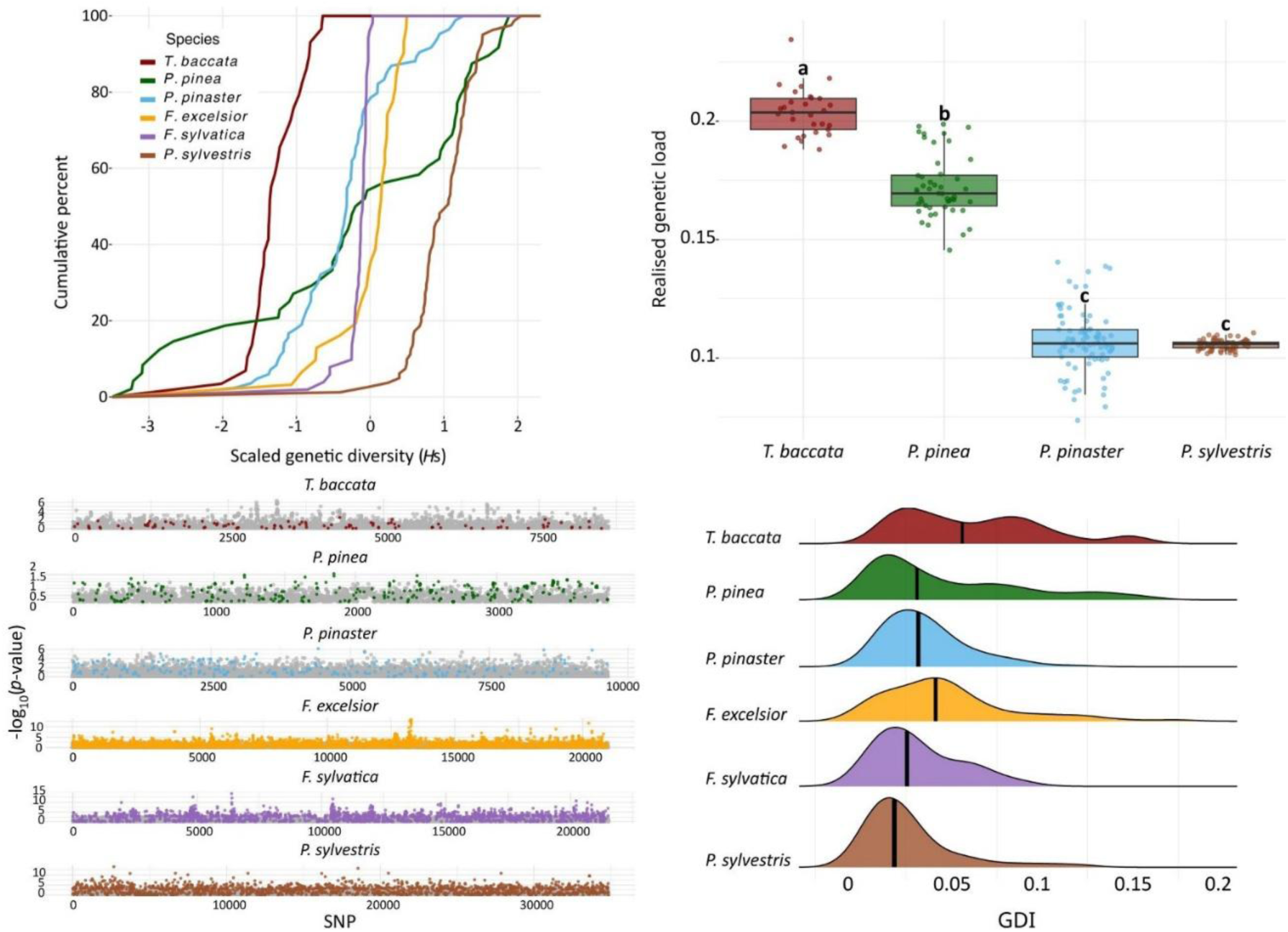
Genetic metrics of population vulnerability to climate change across the studied species. **A** Cumulative distribution of scaled genetic diversity (*H*s; calculated by subtracting the mean and dividing by the standard deviation) across populations. A stepped curve indicates low variability in *H*s across populations (e.g., *F. sylvatica*), whereas a flatter curve indicates high variability (e.g., *P. pinea*). **B** Comparison of realised genetic load across the four conifers species, with Tukey post-hoc groups indicated by lowercase letters. **C** Manhattan plots showing the retained candidate SNPs (coloured) used for GDI calculation, plotted against the total number of SNP, which varied across species (Table S2). *P*-values were obtained from the RDA GEA models. **D** GDI density plot across populations for each species. Black line represents the median GDI values. Note that values are not directly comparable among species.

We also observed marked differences in realised genetic load (i.e., estimated by the number of deleterious mutations in the homozygous state; hereafter referred to as ‘genetic load’; see Methods) among species (Fig. 2B). Interestingly, *T. baccata* and *P. pinea* exhibited the highest genetic load values, whereas *P. pinaster* and *P. sylvestris* had comparable lower medians. Furthermore, among-population variability in genetic load was greatly reduced in *P. sylvestris* compared to the other species (e.g., 0.101 to 0.111 in *P. sylvestris* compared to 0.074 to 0.140 in *P. pinaster*).

Using the best genotype-environment association (GEA) models, we selected candidate single nucleotide polymorphisms (SNPs) to estimate climate adaptation via the genomic discrepancy index (GDI; corresponding to the deviation of a population from the expected GEA models; see Methods), retaining between 100 (*T. baccata*) and 2,957 (*F. excelsior)* SNPs depending on the species (Figs. 2C, S3; Table S2). The distribution of GDI (Fig. 2D) showed considerable variability within and among species. The distributions for *F. sylvatica* and *P. pinaster* were nearly Gaussian, whereas *F. excelsior and P. sylvestris* were still roughly Gaussian but exhibited long tails with higher GDI values. In contrast, *T. baccata* and *P. pinea* displayed very long tails of higher GDI values, approaching a bimodal distribution, with many populations exhibiting elevated GDI. Populations of *T. baccata* with high GDI are primarily found in southern regions, including Greece, Bosnia-Herzegovina, Spain and Italy^17^.

### Population genetic differentiation as a major factor explaining key components of climate change vulnerability

Genetic diversity (*H*s) decreased with increasing population-level genetic differentiation (*F*_ST_) in all species (Table 1), a pattern that was further confirmed by using unbiased Jost’s D (Table S3; see Methods). Associations remained highly significant and consistent across species when atypical population-level *F*_ST_ values were removed and when geography was considered in the models (except for *F. excelsior*; Table 1; Fig. S4; see Methods).

**Table 1.**
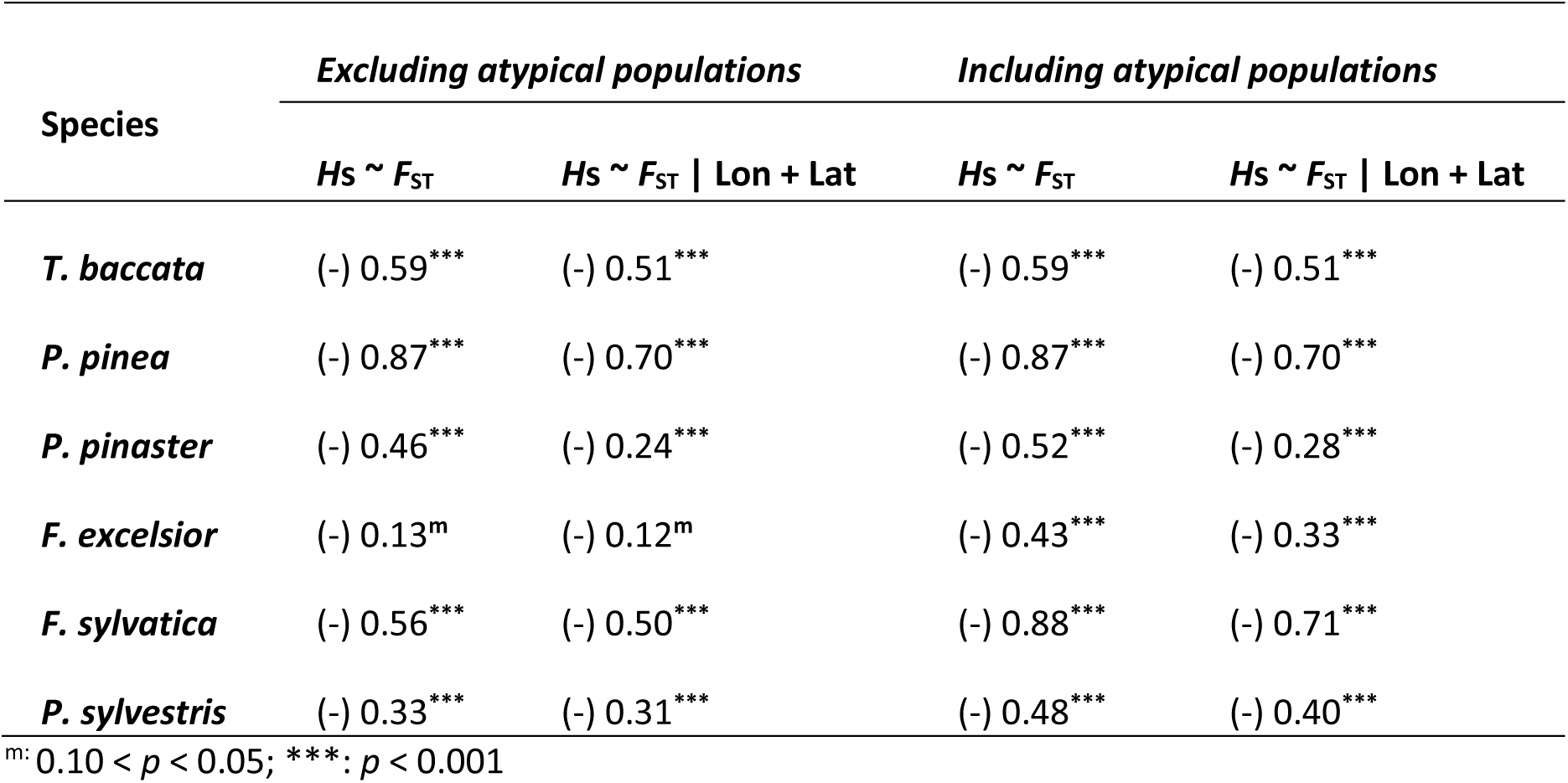
Association between population genetic differentiation (*F*_ST_) and genetic diversity (*H*s). Linear model coefficient signs (β), explained variance (*R*²), and *F*-test *p*-values, summarising the direction and strength of the relationship between population-level *F*_ST_, used here as a proxy for demographic history, and genetic diversity (*H*s). Analyses were run including and excluding populations exhibiting atypically high *F*_ST_ values and accounting or not for geographical variation in *H*s (longitude ‘Lon’ and latitude ‘Lat’).

| Species | Excluding atypical populations |  | Including atypical populations |  |
| --- | --- | --- | --- | --- |
| | $H_s \sim F_{ST}$ | $H_s \sim F_{ST} \mid \text{Lon} + \text{Lat}$ | $H_s \sim F_{ST}$ | $H_s \sim F_{ST} \mid \text{Lon} + \text{Lat}$ |
| *T. baccata* | (-) 0.59*** | (-) 0.51*** | (-) 0.59*** | (-) 0.51*** |
| *P. pinea* | (-) 0.87*** | (-) 0.70*** | (-) 0.87*** | (-) 0.70*** |
| *P. pinaster* | (-) 0.46*** | (-) 0.24*** | (-) 0.52*** | (-) 0.28*** |
| *F. excelsior* | (-) 0.13<sup>m</sup> | (-) 0.12<sup>m</sup> | (-) 0.43*** | (-) 0.33*** |
| *F. sylvatica* | (-) 0.56*** | (-) 0.50*** | (-) 0.88*** | (-) 0.71*** |
| *P. sylvestris* | (-) 0.33*** | (-) 0.31*** | (-) 0.48*** | (-) 0.40*** |

<sup>m</sup>: $0.10 < p < 0.05$ ; \*\*\*: $p < 0.001$

All conifer species, except *P. pinea*, showed a lower genetic load (i.e., a reduction of deleterious mutations in homozygous form) in populations with higher *F*_ST_. In contrast, the increase in genetic load with *F*_ST_ observed in *P. pinea* is likely linked to its unique demographic history, characterised by a relatively recent population collapse^42^. These relationships were consistent whether populations with atypically high population-level *F*_ST_ values were included or not in the analysis (Figs. 3A, S5A). Genetic load was not computed for the two broadleaves (*F. excelsior* and *F. sylvatica*) as its impact is expected to be lower in angiosperm species (see Introduction).

**Figure 3.**
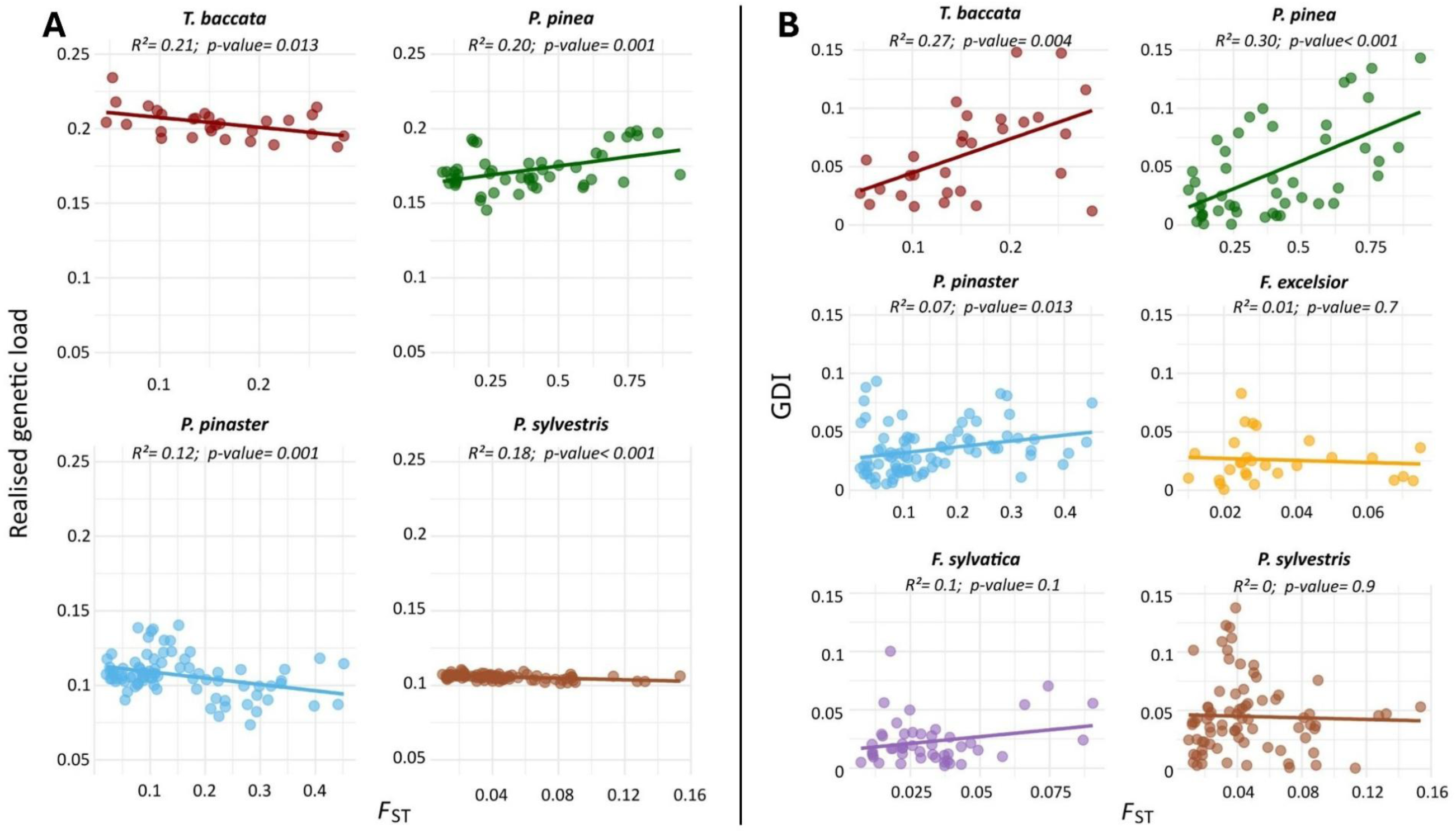
Population-level genetic differentiation (*F*_ST_) as a key factor explaining vulnerability to climate change. Linear relationships between (**A**) realised genetic load (only conifers) or (**B**) GDI, and population-level *F*_ST_, after excluding populations with atypically high *F*_ST_ values (see Methods and Fig. S5 for relationships including atypical populations). The coefficient of determination (*R*²) and *F*-test *p*-values are shown for each model.

Climate adaptation, assessed via GDI, also varied among species in ways that reflected the population size and connectivity gradient illustrated in Fig. 1G, with higher genetic differentiation having from negative to negligible effects on climate adaptation along this gradient (Table 1; Fig. 3B, S5B). In *T. baccata* and *P. pinea*, GDI was higher in populations exhibiting greater *F*_ST_. A similar pattern was observed in *P. pinaster*, *F. excelsior* and *F. sylvatica*, although for the latter two species it was only evident when populations with atypically high population-level *F*_ST_ were included, likely reflecting higher thresholds of isolation required to affect climate adaptation. Finally, no significant association was detected between population-level *F*_ST_ and GDI in the widespread, weakly differentiated *P. sylvestris*.

## Discussion

Our study suggests that, beyond macro-glacial dynamics, local historical events and demography, as reflected in present-day genetic differentiation, explain a substantial portion of the variability in genetic diversity among populations. We consistently observed a reduction in genetic diversity with increasing population-level *F*_ST_. This result remained robust after accounting for geographic effects (Table 1), except for *F. excelsior*. Since geography has previously been linked to glacial-postglacial dynamics^59^, these findings support the additional role of local genetic drift and limited historical gene flow in shaping the observed patterns. Overall, our work suggests that even species capable of long-distance dispersal and therefore substantial gene flow may still experience long-term isolation. This isolation likely led to reduced genetic diversity through a combination of ecological or environmental specialisation, genetic drift, and inbreeding^30,60,61^. Since rapid adaptation mostly relies on standing genetic variation^62^, the general reduction in genetic diversity in highly differentiated populations could limit their adaptive potential, making them particularly vulnerable to climate change^12,39^.

Genetic load varied among species, reflecting differences in their life-history traits and demographic history. A reduction in deleterious mutations with increasing genetic differentiation was observed across all species except *P. pinea*. Historical isolation is expected to intensify genetic drift, reducing effective population size and increasing inbreeding. This can increase the fixation of deleterious alleles, thereby strengthening purifying selection and ultimately reducing realised genetic load^63,65^. Similar patterns have been reported in other long-lived organisms, where increased genetic drift and isolation have led to the purging of deleterious mutations, likely increasing their adaptive potential^37,66^. The unique pattern observed in *P. pinea* (positive association with genetic differentiation) can also be explained by demographic history. Present-day gene pools of this species likely persisted through near-extinction in a small number of Iberian populations with fossil records dating back to the last Neanderthals (c. 49 kyr BP)^42,67^. Under such prolonged and extreme isolation, reduced effective population size and limited gene flow likely promoted the accumulation, rather than the purging, of deleterious mutations^65,68^.

Across species, genetic load tracked gradients of population size and connectivity (Fig. 1G), with higher values observed in *T. baccata* and *P. pinea* than in *P. pinaster* and *P. sylvestris*. This pattern suggests that species with smaller and more fragmented populations tend to accumulate more deleterious mutations, regardless of within-species dynamics. Consistent with this, demographic history and the resulting patterns of population differentiation account for much of the variation in genetic load across conifer populations. In particular, the accumulation of deleterious mutations appears to be shaped by historical effective population size and connectivity, underscoring the importance of considering these factors when assessing the consequences of genetic load for population persistence under climate change.

Increasing genetic differentiation was associated with suboptimal climate adaptation in all species except *P. sylvestris*. Alongside the parallel reduction in genetic diversity with increasing differentiation, these results highlight the potential beneficial role of gene flow in enhancing adaptation by maintaining standing genetic variation, a key driver of adaptive evolution^33,69^. Furthermore, both theoretical predictions and empirical evidence also showed that, by introducing pre-adapted alleles, gene flow can further facilitate climate tracking, particularly in leading-edge populations^32,70,71^. We observed marked interspecific differences in climate adaptation in our study. Species with a prevalence of small and fragmented populations (e.g., *T. baccata* and *P. pinea*) showed stronger negative associations between population genetic differentiation and climate adaptation than species with larger, more continuous populations (e.g., *F. sylvatica* and *P. sylvestris*). These differences may, at least in part, reflect variation in historical effective population size, as the benefits of increased standing genetic diversity and enhanced climate tracking via gene flow are likely to be more pronounced in smaller populations^32,71,72^.

Interestingly, across the six tree species, populations with low genetic differentiation consistently showed no signs of reduced climate adaptation. These results provide further empirical support for the positive role of genetic connectivity in enhancing climate adaptation in small, fragmented populations, while showing no detectable negative effects in larger, more continuous populations. However, while biologically meaningful, these insights need to be taken with caution. For instance, climate adaptation models used here rely on several assumptions, including that genotype-environment associations accurately capture adaptive signals^73^. Furthermore, higher Euclidean distances between observed and predicted genomic compositions (GDI) were interpreted here as signatures of suboptimal climate adaptation, therefore neglecting other potential explanations, such as alternative adaptive pathways or pre-adaptation to future climate.

### Conclusion

Genomic analyses of 6,435 trees from 326 populations across six ecologically and economically important forest tree species revealed major effects of demographic history, as summarised by population-level genetic differentiation, on shaping population vulnerability to climate change. Population adaptive potential, reflected in within-population genetic diversity, declined consistently with historical isolation. At the same time, sensitivity to climate change was likely shaped by reduced effective population size and limited historical gene flow, leading either to the purging or the accumulation of deleterious mutations, and increasing maladaptation to contemporary climates. Overall, this work provides valuable insights into how genetic differentiation, reflecting genetic drift and limited historical gene flow, influences current vulnerability of forest tree populations to climate change. Future empirical work could aim to link genetic metrics to effective population size, phenotypic performance, or reproductive fitness, to better assess the contribution of each factor to climate vulnerability. Our findings can also inform management strategies, including assisted gene flow, by helping to identify populations most at risk from climate change and those likely to benefit from increased genetic variance in fitness-related traits^74^. Assessing current gene flow dynamics will be crucial to determine whether rapid environmental change is already reshaping historical patterns of connectivity and genetic differentiation.

## Materials and methods

### Sampling and genotyping

Our study focused on six European tree species, namely *Taxus baccata*, *Pinus pinea*, *Pinus pinaster*, *Fraxinus excelsior, Fagus sylvatica* and *Pinus sylvestris.* Species were selected to represent contrasting demographic histories, including different phylogenetic groups, stand formation and population sizes, as well as levels of connectivity and population genetic structure^30,43^ (Fig. 1G; see Results). For each species, 29-86 populations spanning most of their ranges were sampled, with total sample sizes per species ranging from 490 to 1,729 mature trees (except *for F. excelsior* where young trees were sampled; see Supplementary Files 1 and 2). Particular attention was paid to sample populations with different degrees of genetic differentiation, including both core and range-edge populations.

Genomic data for 490 trees representing 29 populations of *T. baccata* were obtained from a previous study^17^. This dataset comprised single nucleotide polymorphisms (SNPs) generated by combining gene-capture and single-primer enrichment technology (SPET) data.

Genomic data for 461 trees from 31 natural populations of *F. excelsior* were compiled from one previous data paper and one study: 286 trees sequenced using SPET^75^ and 175 using low-coverage whole genome sequencing^76^, following Doonan *et al*.^77^. SNPs were called against the same reference genome, FRAX_0001_PL^78^.

Genomic data for 1,224 trees from 51 populations of *F. sylvatica* were generated by combining two previous datasets: 370 trees from a data paper^75^ and 854 trees from a published dataset^79^, both of which produced using SPET. SNP quality filtering and the removal of outliers and duplicates were performed on the merged dataset following the procedure described in Pinosio *et al*.^75^.

Genomic data for the three pine species (*P. pinea*: 952 trees from 48 populations; *P. pinaster*: 1,579 trees from 86 populations; *P. sylvestris*: 1,729 trees from 81 populations) were retrieved: from a previous study^80^ for *P. pinaster* and from a data paper^81^ for *P. pinea* and *P. sylvestris*. SNP genotypes were generated using the multispecies 4TREE (*P. pinea*, *P. pinaster*) and the PiSy50k (*P. sylvestris*) Axiom arrays (Thermo Fisher Scientific, USA) at IGA Technology Services (Udine, Italy). Following Olsson *et al*.^81^, we applied the quality thresholds for markers CR ≥ 85, FLD ≥ 3.2, HetSO ≥ −0.3 and samples DQC ≥ 0.4 with QC call rate ≥ 85, in order to maximise SNP retention while ensuring high data quality.

Overall, genomic datasets included 6,435 trees from 326 populations, mainly in Europe and bordering regions in North Africa and Asia (see details in Supplementary Files 1 and 2). For each species, we performed population genetic structure analyses to ensure that merged datasets were biologically consistent and appropriate for downstream analyses (Figs. S6, S7). Two of the 326 populations (COM and FUEbis of *P. pinaster*) were excluded because they were duplicated or had too few samples. For each species, two datasets were then generated after removing monomorphic SNPs and variants that failed additional filtering thresholds (coverage depth < 7 or > 150, and allele balance at heterozygous sites < 0.2 or > 0.8). The first dataset (*dataset 1*) retained SNPs and individuals with less than 15% missing data and SNPs with a minor allele count of at least three (except for *T. baccata*, see Francisco *et al*.^17^). This dataset was used for population genetic structure (PCA, STRUCTURE) and differentiation (population-level *F*_ST_) analyses, as well as for calculating genetic diversity (*H*s), as these analyses are sensitive to high rates of missing data and low-frequency SNPs^82^. *Dataset 1* comprised between 3,563 (*P. pinea*) and 34,760 SNPs (*P. sylvestris*), depending on the species, with a total of 6,262 trees genotyped across the 324 populations (Table S4). The second dataset (*dataset 2*), used for computing climate adaptation and genetic load statistics, was filtered out for missing data > 30% (both for SNPs and individuals) and for minor allele counts lower than twice the number of individuals in the smallest population before filtering (Table S4). This latter filter was applied to remove low-frequency alleles that could otherwise increase false-positive rates, while retaining potential climate-associated SNPs or deleterious mutations that are fixed in small populations. *Dataset 2* included between 3,790 (*P. pinea*) and 34,848 SNPs (*P. sylvestris*), depending on the species, with a total of 6,326 trees genotyped across the 324 populations (Table S4). From these datasets, SNPs from coding regions that could be polarised and annotated using the *Pinus tabuliformis* or *Taxus chinensis* reference genomes^83,84^ were used to estimate genetic load (see below). This yielded 4,700; 1,195; 2,219; and 12,808 SNPs for *T. baccata*, *P. pinea*, *P. pinaster*, and *P. sylvestris*, respectively. Finally, for climate adaptation analyses, missing data in *dataset 2* were imputed using the most common genotype within each gene pool.

### Climatic data

Climatic data were extracted at high spatial resolution (30 arc-seconds) from Climate Downscaling Tool (ClimateDT^85^). The 1901-1950 period was defined as the reference climate to which the populations are currently adapted, except for *F. excelsior*, for which the 1991-2020 period was used instead. This more recent time interval may better reflect the climate to which populations are adapted in this species, given the relatively young age of the sampled trees and the species’ known capacity for rapid adaptation^86^. For each species, the retained climatic variables (Table S5) were selected following the approach described in Francisco *et al*.^17^: (i) a pre-selection step based on species life-history traits and the available literature, (ii) forward selection analysis based on genomic importance, and (iii) removal of correlated variables (Pearson’s |*r*| > 0.75) to ensure that multicollinearity remained low and the variance inflation factor below 10 across variables (see Supplementary Material).

### Population genetic structure and differentiation

For each species, principal component analysis (PCA) was used to assess population genetic structure, as implemented in the VEGAN R package^87^ (Fig. S6). Genetic clusters were also examined based on previously^17,88^ or newly run Bayesian clustering analyses, as implemented in STRUCTURE v2.3.4^89^ (Fig. S7). For newly run STRUCTURE analyses, ten independent MCMC runs were performed for each number of potential clusters (*K* = 1-10), with 100,000 iterations per run and a burn-in of 10,000 iterations. The most probable number of genetic clusters was determined using the ΔK method^90^.

Population-level genetic differentiation (*F*_ST_), used here as a proxy for population-specific demographic history, encompassing historical population size fluctuation, gene flow and connectivity (e.g.,^91,92^), was calculated using BayeScan v2.1^93^. In this approach, *F*_ST_ represents the genetic differentiation of each population relative to a predicted ancestral gene pool, making it less influenced by local outliers and more reflective of broad historical patterns of divergence. Given the comprehensive sampling effort, including both core and range-edge populations for each species, populations with high *F*_ST_ can be considered more historically isolated than the species average.

### Genetic metrics of vulnerability to climate change

Genetic differences among individuals of the same species can produce diverse phenotypes within populations, some of which may confer advantages in shifting climate. The extent of this genetic diversity can determine the population’s evolutionary potential^12^. Genetic variation within populations was quantified using Nei’s genetic diversity (*H*s), as implemented in the HIERFSTAT R package^94^. Due to the differences in experimental design to obtain genomic data (e.g., gene capture, SPET designs and different genotyping arrays), genetic diversity values cannot be compared across species.

Genetic load was estimated only for the conifer species in this study (*T. baccata*, *P. pinea*, *P. pinaster*, and *P. sylvestris*), as the accumulation of deleterious mutations in these species has been shown to be greater than in broadleaves and linked to fitness-related traits (e.g.,^95^; see Introduction). First, SNPs were mapped onto the *Pinus tabuliformis* reference genome^83^ (*P. pinea*, *P. pinaster* and *P. sylvestris*; see Supplementary Material) or onto the *Taxus chinensis* reference genome^84^ (*T. baccata*). SnpEff software (version 5.1) was then used to identify SNPs causing amino acid changes in protein-coding sequences^96^. Finally, the functional impact of mutations was predicted using PROVEAN (version 1.1.5), a software that assesses the impact of mutations by evaluating sequence conservation and alignment scores across homologous proteins^97^. For each SNP, PROVEAN predicted its impact on the biological function of the corresponding protein and assigned a ‘deleteriousness’ score. Mutations were considered deleterious when the PROVEAN score was lower than −2.5^97^. Realised genetic load was then computed following González-Martínez *et al*.^27^, as the ratio of the expected number of deleterious mutations in homozygosis to the number of expected neutral mutations. As most deleterious alleles are thought to be recessive, the realised genetic load is therefore expected to have a stronger immediate impact on fitness than the absolute number of deleterious mutations^27,65^.

Population climate adaptation was quantified using a two-step procedure. First, we investigated the presence of genomic markers potentially involved in climate adaptation by carrying out genotype-environment association (GEA) analyses. For each species, six GEA methods (correcting or not for population genetic structure) were applied to identify candidate loci: redundancy analysis (RDA), partial redundancy analysis (pRDA), latent factor mixed models (LFMM), BayPass, gradient forest (GF) uncorrected for population genetic structure (GF-raw) and GF corrected for population genetic structure (GF-corrected; see Table S6 and Supplementary Material for details). We then defined an *all-outlier* SNP set comprising all candidate SNPs identified across the GEA analyses. Following Archambeau *et al*.^98^, an *outlier* SNP set was further defined, including only unlinked candidate SNPs identified by at least two GEA methods. For this, we used a linkage disequilibrium threshold of *R*² < |0.7| within contigs or 2,000 bp, depending on the available information for each species, and filtered the datasets for linked loci using the GENETICS R package^99^. For each species, a *random* SNP set was also generated by randomly selecting SNPs not identified as candidates by any GEA method, matching both the number and allele frequency distribution of the *outlier* set (see Supplementary Material).

Second, climate adaptation was quantified for each population within species by computing the genomic discrepancy index (GDI^17^) using the RDA method. GDI computes the Euclidean distances between the observed and predicted population scores along the retained RDA axes. Higher population GDI indicates a greater deviation from the established GEA relationship relative to other populations. For most species, the *all-outlier* SNP set was used to build the reference GEA model for GDI estimates. However, for *T. baccata* and *P. pinaster*, we applied SNP sets (*random* and *outlier*, respectively) previously identified as most suitable through model evaluation with fitness proxies^17,98^.

For all species, GDI estimates based on alternative SNP sets were highly correlated with those obtained from the selected sets (Pearson’s correlation > 0.64; Table S2).

### Linking demographic history to vulnerability to climate change

To assess the effects of demographic history on population vulnerability to climate change, we examined, for each species, the relationship between population-level *F*_ST_ and genetic diversity, genetic load and GDI (see Supplementary File 3). Associations were evaluated using linear models with *F*-test *p*-values, as implemented in the STATS R package^100^. To avoid interpreting associations driven solely by extreme values, linear models were investigated with and without populations exhibiting atypically high *F*_ST_, defined as those falling outside 2.4 times the interquartile range of population-level *F*_ST_ within each species.

## Author Contributions

Designed research: TF, AP, SCG-M; acquired and pretreated genomic data: SO, MK, MW, SP, GGV, KBB, LRN, JMD, DG, EV; analysed data: TF, IL-K, SCG-M; wrote the paper: TF, SCG-M; obtained funding: MW, GGV, DG, AP, SCG-M, LRN; all authors participated in discussions and revised multiple versions of the manuscript, including the final version.

## Acknowledgments

TF was funded by a PhD grant of the INRAE Scientific Department ECODIV. EV was supported by the MSCA European fellowship MedForAct (GA 101107604). MK was supported by the Slovenian Research Agency (research core funding no. P4-0107 and Young Researcher grant no. 58170). This research was funded by EU H2020 FORGENIUS (grant agreement No. 862221) and Horizon Europe OptFORESTS (grant agreement No. 101081774) projects. Views and opinions expressed are however those of the authors only and do not necessarily reflect those of the European Union. Neither the European Union nor the granting authority can be held responsible for them. Additional *F. excelsior* data was generated by the AshAdapt project supported by the Independent Research Fund Denmark (DFF Technology and Production Sciences) under grant no. 8022-00355B. We are grateful to the Genotoul bioinformatics platform Toulouse Occitanie (Bioinfo Genotoul, https://bioinfo.genotoul.fr/) for providing computing and storage resources.

## Data Archiving Statement

Genomic data for *Taxus baccata* from Francisco *et al*.^17^ is available at Data INRAE (doi: https://doi.org/10.57745/JJ0ZEI); for *Pinus pinaster* from Olsson *et al*.^80^ at Zenodo (doi: 10.5281/zenodo.14950394); for *Pinus pinea* and *Pinus sylvestris* from Olsson *et al*.^81^ at Zenodo (doi: 10.5281/zenodo.17455887 and 10.5281/zenodo.17462035, respectively). Newly combined genomic data for *Fraxinus excelsior* and *Fagus sylvatica* are openly available at Data INRAE (doi: 10.57745/OG2FBY).

## Code availability

Code used for data processing and analysis will be made publicly available upon publication at a GitHub public repository.

