## Supplementary information for "Demographic history shapes forest tree vulnerability to climate change"

### *Supplementary Methods*

#### *Climatic variable selection*

To identify the climatic variables potentially driving adaptation in each selected species, we implemented a three-step procedure (see Main text). In the second step, forward selection was applied to the preselected variables (from step one) to identify those explaining the largest proportion of genomic variation among populations. This was achieved using the ordiR2step function from the VEGAN R package[1], by evaluating models ranging from a null model with no explanatory variables up to a full model incorporating all candidate variables. Variables were retained based on their contribution to increasing the model's $R^2$, with significance assessed via 1,000 permutations and a *p-value* threshold of 0.05; twenty runs were performed to ensure robustness. In the third step, climatic variables showing high correlation (Pearson's $|r| > 0.75$) and/or variance inflation factors (VIF) over 10 (see James *et al.*[2]) were excluded from the analyses, prioritising those identified as the most informative in the forward selection step. The resulting set of climatic variables contained either six or seven variables depending on the species (Table S5).

#### *Genetic load annotation for pines*

For the three pine species, SNPs were mapped onto the *Pinus tabuliformis* reference genome[3]. Due to the large size of its chromosomes, a working version of the reference genome was first generated by splitting the 12 chromosomes into 22,739 contigs[4]. SNPs from the three pine species were then transferred onto the *P. tabuliformis* genome based on the sequence similarity of their 120 bp flanking regions using BLASTN (NCBI BLAST 2.2.26). For each SNP, the best alignment was retained based on percentage identity, alignment length, number of mismatches, and e-value.

#### *Climate-related candidate SNPs and other SNP sets*

Several gene-environment associations (GEA) methods were applied to each species to identify climate-related candidate SNPs (see below and Table S6).

RDA and pRDA are linear multivariate canonical analyses that combine ordination and multiple regression approaches to produce canonical axes[5]. In the pRDA, genetic structure was accounted for using the first two axes of a principal component analysis based on the SNP *dataset 1* (see Table S4 and Main text). Following Capblancq and Forester[6], SNPs with an extreme Mahalanobis distance from the centre of the RDA space along the significant axes were considered as candidate loci.

Latent Factor Mixed Model (LFMM) is a multivariate linear regression method. It uses least squares estimation to assess the effect size of each locus[7], and latent factors to account for population genetic structure.

BayPass standard covariate (STD) model is a Bayesian univariate method that estimates the extent to which each marker is linearly associated with the predictors[8]. A variance-covariance matrix computed from the SNP *dataset 1* (see Table S4 and Main text) was used to account for population genetic structure. SNPs with a mean Bayes factor (BF) greater than the species threshold across five independent runs, corresponding to either 'substantial evidence' or 'very strong evidence' on the Jeffreys' scale[9], were considered candidate loci for climate adaptation.

Gradient Forest (GF) is a non-linear machine learning algorithm that fits an ensemble of regression trees in order to model changes in allele frequencies across populations[10,11]. GF models were applied to either raw allele frequencies (GF-raw) or allele frequencies corrected for population genetic structure, obtained by multiplying the LFMM-adjusted effect size matrix by the transpose of the climatic variable matrix (GF-corrected)[12]. For each model, five independent runs were performed, retaining the top 1% or 5% of SNPs, depending on the species. Then, the overlapping SNPs across the five runs were considered candidate loci.

To generate the *outlier* SNP set (see Main text), two thresholds were used to identify candidate SNPs with the GEA methods described above. To ensure that GDI models were based on a sufficient number of SNPs, thresholds were adjusted across species. For *T. baccata*, *P. pinea*, and *F. excelsior*, the less conservative set of thresholds was applied, whereas for *P. pinaster*, *F. sylvatica*, and *P. sylvestris*, the more conservative one was used (see Table S2 and S6 for details).

### *Supplementary Tables & Figures*

*Table S1.* **Association between genetic diversity (*Hs*) and geographical variables**. Linear model coefficient signs (β), explained variance ($R^2$), and *F*-test *p*-values, summarising the direction and strength of the relationship between genetic diversity (*Hs*) and geographical coordinates (longitude and latitude).

| **Species** | ***Hs* ~ Longitude** | ***Hs* ~ Latitude** |
|---|---|---|
| ***Taxus baccata*** | (+) 0.21* | $0.07^{ns}$ |
| ***Pinus pinea*** | (-) 0.14** | $0.00^{ns}$ |
| ***Pinus pinaster*** | (-) 0.12** | (+) 0.06* |
| ***Fraxinus excelsior*** | $0.00^{ns}$ | $0.01^{ns}$ |
| ***Fagus sylvatica*** | (+) 0.20** | $0.00^{ns}$ |
| ***Pinus sylvestris*** | (+) 0.19*** | (+) 0.18*** |

$^{ns}$: not significant; *: $p < 0.05$; **: $p < 0.01$; ***: $p < 0.001$

*Table S2*. **Climate-related candidate SNP sets**. Number of climate-related candidate loci identified using GEA, along with the number of candidate loci retained in each species and the Pearson's correlation of GDI values across the three SNP sets (see Main text).

| **Species** | **Number candidate SNPs (all/retained)** | **Selected SNP set (threshold for candidate identification)** | **Mean Pearson's correlation of GDI across SNP sets** |
|---|---|---|---|
| ***T. baccata*** | 935/100 | *random* (LC) | 0.78 |
| ***P. pinea*** | 296/296 | *all-outlier* (LC) | 0.64 |
| ***P. pinaster*** | 1178/253 | *outlier* (MC) | 0.71 |
| ***F. excelsior*** | 2957/2957 | *all-outlier* (LC) | 0.92 |
| ***F. sylvatica*** | 2085/2085 | *all-outlier* (MC) | 0.83 |
| ***P. sylvestris*** | 1595/1595 | *all-outlier* (MC) | 0.97 |

LC: less conservative; MC: more conservative; see Table S6.

*Table S3*. **Correlation between population-level $F_{ST}$ and Jost's D**. Pearson's correlation between population-level $F_{ST}$, as estimated by BayeScan, and Jost's D, a genetic divergence estimator that is not correlated by construction with within-population genetic diversity.

| Species | Pearson's correlation |
|---|---|
| ***T. baccata*** | 0.93 |
| ***P. pinea*** | 0.94 |
| ***P. pinaster*** | 0.97 |
| ***F. excelsior*** | 0.96 |
| ***F. sylvatica*** | 0.99 |
| ***P. sylvestris*** | 0.91 |

*Table S4*. **Genomic datasets**. Description of the genomic datasets used in the various analyses. Minor allele count (MAC) thresholds in *dataset 2* correspond to twice the number of individuals in the smallest population prior to filtering (see Methods for details on the filtering steps).

| **Species** | **Population genetic structure, genetic differentiation and genetic diversity (*dataset 1*)** | **Climate adaptation and genetic load (*dataset 2*)** | |
|---|---|---|---|
| | | **MAC threshold** | **Dataset** |
| ***T. baccata*** | 452 trees; 29 populations; 8,252 SNPs | 20 | 475 trees; 29 populations; 8,616 SNPs |
| ***P. pinea*** | 946 trees; 48 populations; 3,563 SNPs | 24 | 952 trees; 48 populations; 3,790 SNPs |
| ***P. pinaster*** | 1,559 trees; 84 populations; 8,684 SNPs | 12 | 1,551 trees; 84 populations; 9,636 SNPs |
| ***F. excelsior*** | 418 trees; 31 populations; 21,037 SNPs | 14 | 443 trees; 31 populations; 20,997 SNPs |
| ***F. sylvatica*** | 1,160 trees; 51 populations; 30,076 SNPs | 46 | 1,177 trees; 51 populations; 21,529 SNPs |
| ***P. sylvestris*** | 1,727 trees; 81 populations; 34,760 SNPs | 16 | 1,728 trees; 81 populations; 34,848 SNPs |

*Table S5.* **Climatic variables**. Climatic variables retained in the genotype-environment association (GEA) models for each species. Bio1: Annual mean temperature; Bio2: Mean diurnal range; Bio3: Isothermality; Bio4: Temperature seasonality; Bio6: Minimum temperature of the coldest month; Bio9: Mean temperature of the driest quarter; Bio10: Mean temperature of the warmest quarter; Bio11: Mean temperature of the coldest quarter; Bio12: Annual precipitation; Bio14: Precipitation of the driest month; Bio15: Precipitation seasonality; Bio18: Precipitation of the warmest quarter; AHM: Annual heat moisture index; SHM: Summer heat moisture index.

| Variables | *Taxus baccata* | *Pinus pinea* | *Pinus pinaster* | *Fraxinus excelsior* | *Fagus sylvatica* | *Pinus sylvestris* |
|---|---|---|---|---|---|---|
| ***Bio1*** | ■ | ■ | ■ | | ■ | |
| ***Bio2*** | ■ | | | ■ | ■ | |
| ***Bio3*** | | ■ | ■ | ■ | | ■ |
| ***Bio4*** | ■ | ■ | ■ | | ■ | |
| ***Bio6*** | | | | ■ | | |
| ***Bio9*** | ■ | ■ | | | ■ | |
| ***Bio10*** | | | | | | ■ |
| ***Bio11*** | | | | | | ■ |
| ***Bio12*** | ■ | ■ | ■ | | ■ | ■ |
| ***Bio14*** | | ■ | | | | |
| ***Bio15*** | ■ | ■ | ■ | ■ | ■ | ■ |
| ***Bio18*** | | | | ■ | | ■ |
| ***AHM*** | | | | ■ | | ■ |
| ***SHM*** | | | ■ | | ■ | |

*Table S6.* **Detection of climate-related candidate SNPs**. Description of the six genotype-environment association (GEA) analyses used to identify climate-associated loci (adapted from Francisco *et al.*[12]). MC: more conservative threshold, applied to *P. pinaster, F. sylvatica*, and *P. sylvestris*; LC: less conservative threshold, applied to *T. baccata, P. pinea*, and *F. excelsior* (see Methods).

| **Method** | **Functional shape** | **Population genetic structure correction** | **Procedure for candidate-SNP selection** | **References** |
| --- | --- | --- | --- | --- |
| Redundancy analysis (RDA) | Linear | No | Extreme Mahalanobis distance with false discovery rate of 5% (MC) or 10% (LC) | Legendre and Legendre[5]; Capblancq and Forester[6] |
| Partial redundancy analysis (pRDA) | | Yes, with two principal component axes | | |
| Latent factor mixed model (LFMM) | Linear | Yes, with two latent factors | False discovery rate of 5% (MC) or 10% (LC) | Caye *et al.*[7] |
| BayPass | Linear | Yes, with covariance matrix | Mean Bayes factor across five runs > 8 (LC) or 10 | Gautier[8] |

| | | | | |
|---|---|---|---|---|
| | | | (MC; 'substantial evidence' or 'very strong evidence' on the Jeffreys' scale; Kass & Raftery[9] | |
| Gradient Forest (GF-raw) | Non-linear | No | Top 1% (MC) or 5% (LC) overlapping SNPs across five runs | Ellis *et al.*[10]; Fitzpatrick *et al.*[11] |
| Gradient Forest corrected (GF-corrected) | | Yes, with LFMM corrected matrix | | |

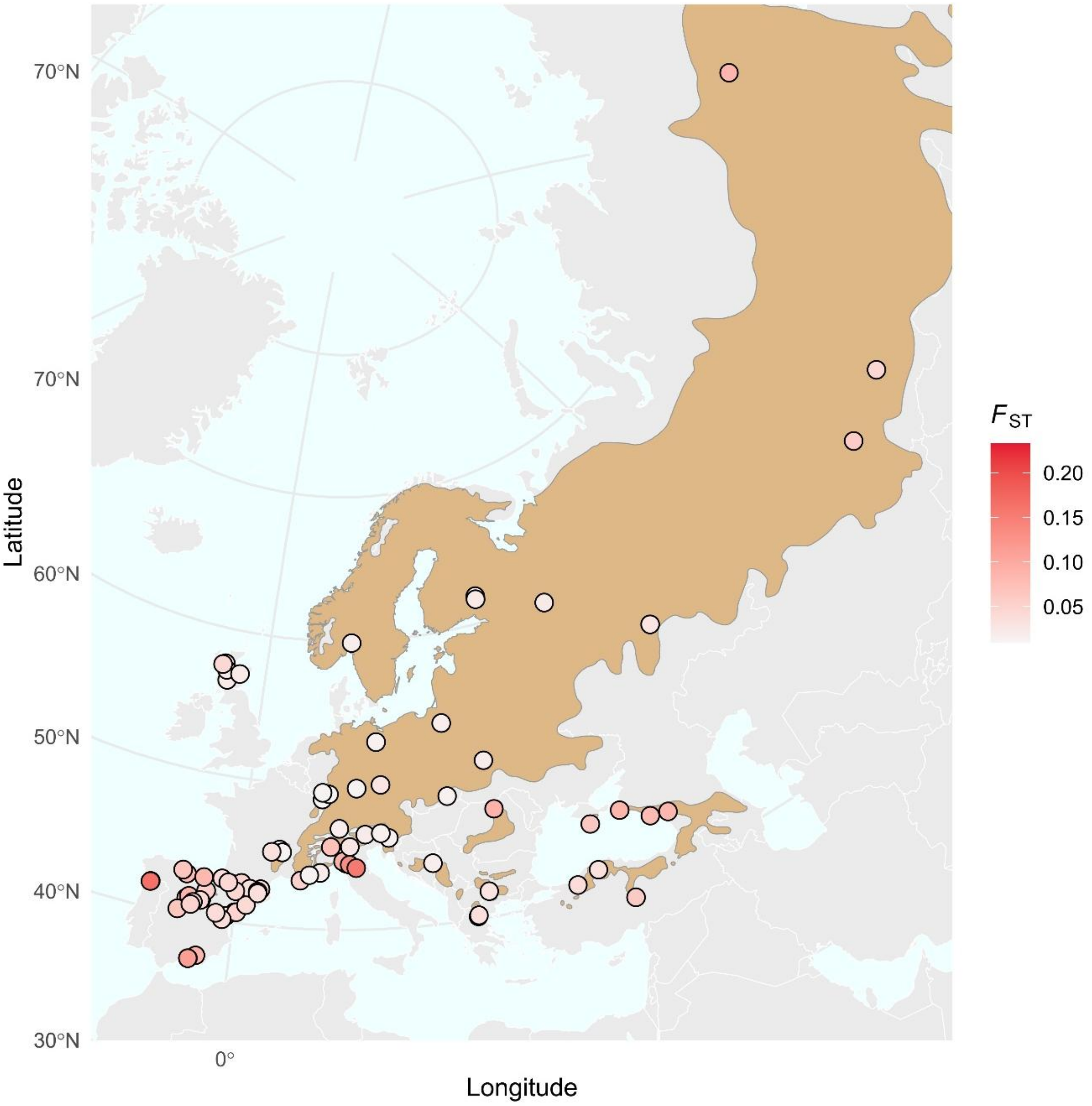


*Figure S1.* **Population-level genetic differentiation (*$F_{ST}$*) across the entire sampled range of *P. sylvestris***. Population-level genetic differentiation ($F_{ST}$) across all sampled populations. Each circle represents a population, with colour intensity indicating the magnitude of $F_{ST}$. Shaded areas depict the approximate species range, as extracted from Caudullo *et al.*[13].

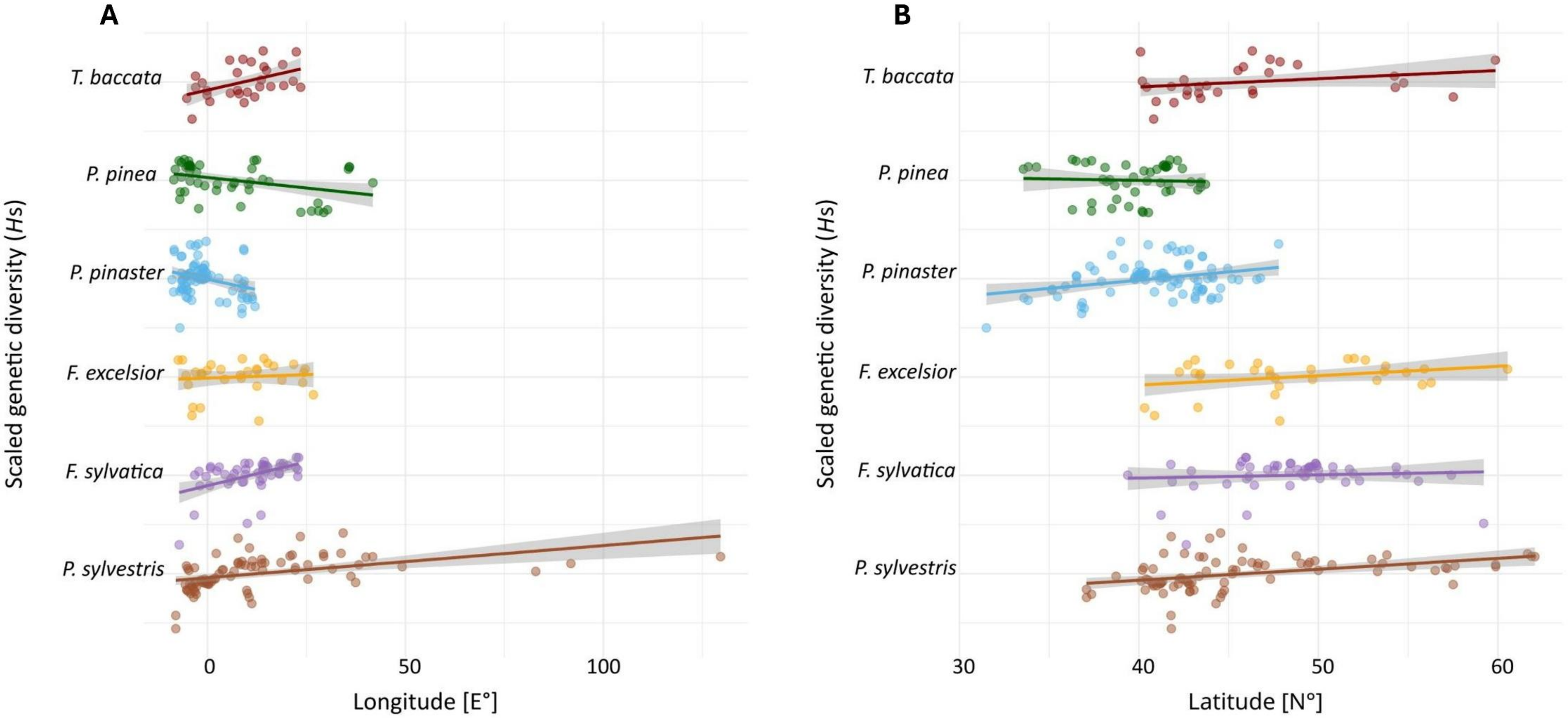


*Figure S2.* **Genetic diversity (*Hs*) trends along longitudinal and latitudinal gradients**. Linear relationships between Nei's genetic diversity (*Hs*) and geographical coordinates. **A** The relationship with longitude was positive for *T. baccata*, *F. sylvatica* and *P. sylvestris*, negative for *P. pinea* and *P. pinaster*, and non-significant for *F. excelsior*. **B** Along latitude, only *P. pinaster* and *P. sylvestris* exhibited a significant (positive) relationship (see also Table S1)

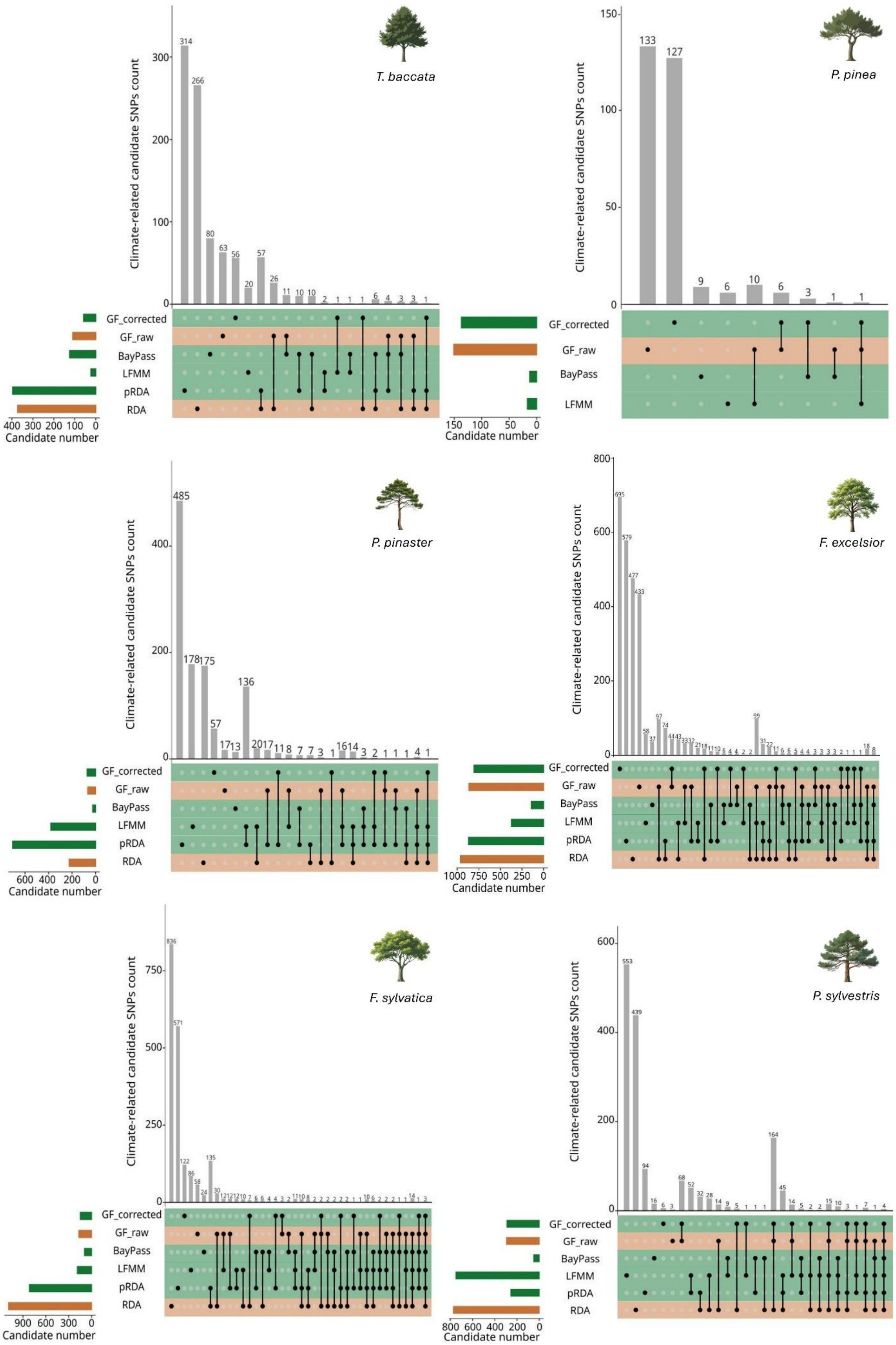
T. baccata
P. pinea
P. pinaster
F. excelsior
F. sylvatica
P. sylvestris
Climate-related candidate SNPs count
Candidate number
GF_corrected
GF_raw
BayPass
LFMM
pRDA
RDA

*Figure S3.* **Climate-associated candidate SNPs as identified by multiple GEA methods**. UpSet plot showing climate-related SNPs per method and across species. Horizontal bars represent the number of candidate SNPs identified by each GEA method, while vertical bars indicate the number of candidate SNPs unique to a single method or shared across multiple methods, with the corresponding method(s) indicated by the dots directly below the vertical bars. Colours refer to whether the method corrects (in green) or not (in orange) for population genetic structure. No candidate markers were identified by RDA and pRDA for *P. pinea*.

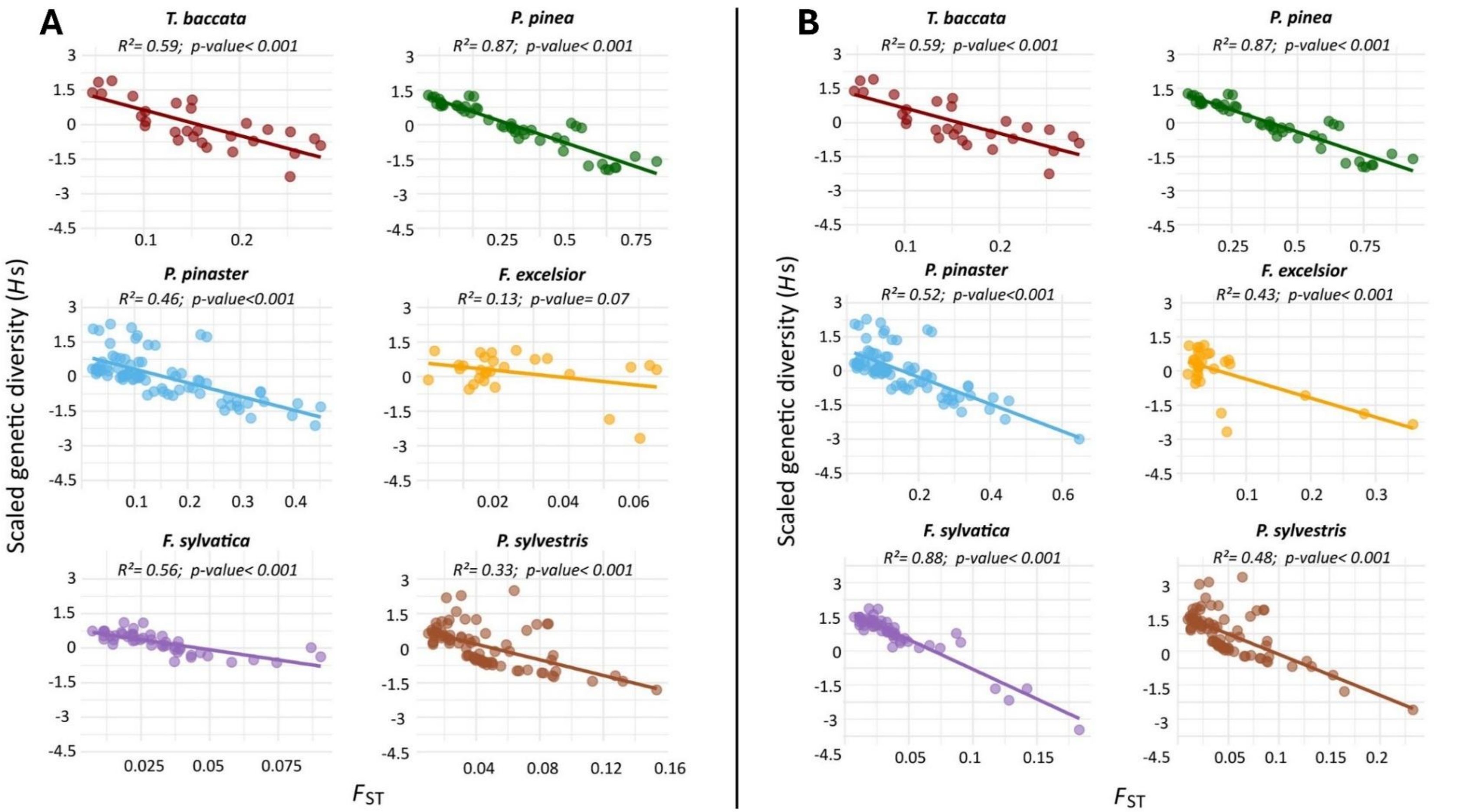


*Figure S4.* **Population-level genetic differentiation ($F_{ST}$) as a driver of within-population genetic diversity.** Linear relationships between Nei's genetic diversity (*Hs*) and population-level $F_{ST}$, (**A**) without or (**B**) with populations exhibiting atypically high $F_{ST}$ values. The coefficient of determination ($R^2$) and *F*-test *p*-values are shown for each model.

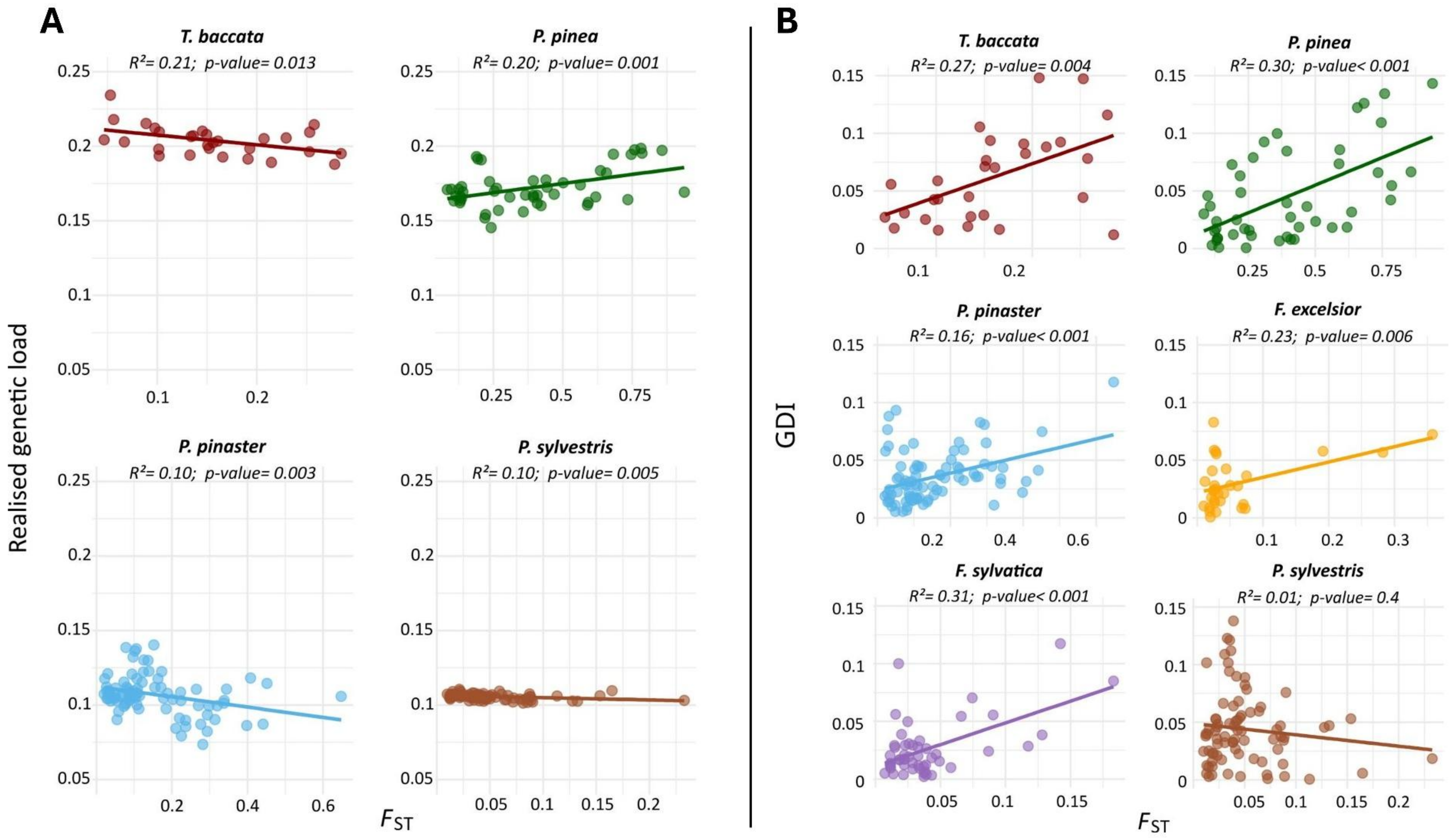


*Figure S5.* **Population-level genetic differentiation ($F_{ST}$) as a key factor explaining sensitivity to climate change.** Linear relationships between (**A**) realised genetic load or (**B**) GDI, and population-level $F_{ST}$, including populations with atypically high $F_{ST}$. The coefficient of determination ($R^2$) and *F*-test *p*-values are shown for each model. Panel (A) only includes conifer species.

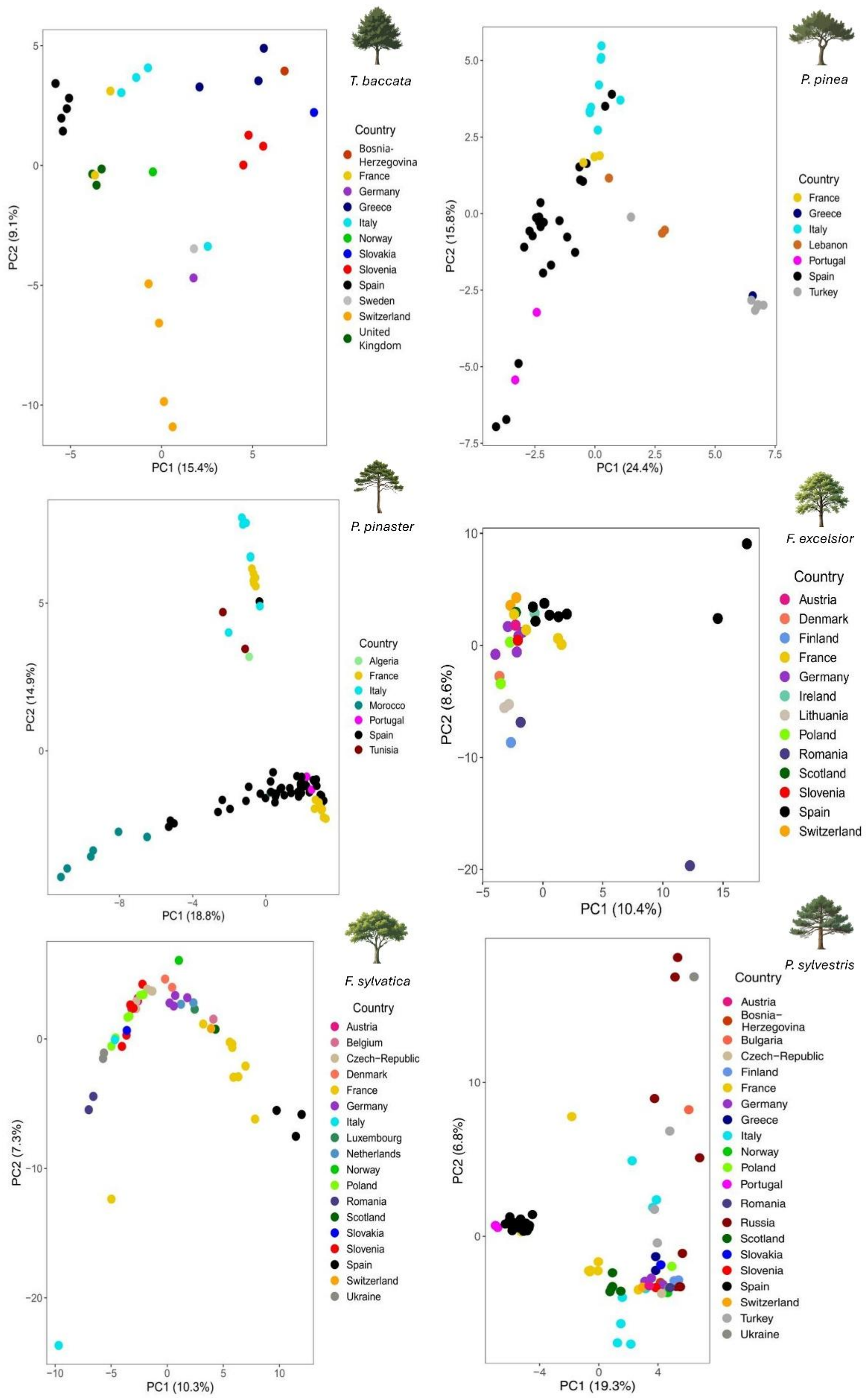


*Figure S6*. **Population genetic structure across the six species**. Principal component analysis (PCA) at the population level, showing the position of each population on the first two axes of the PCA. Colours refer to the country of origin.

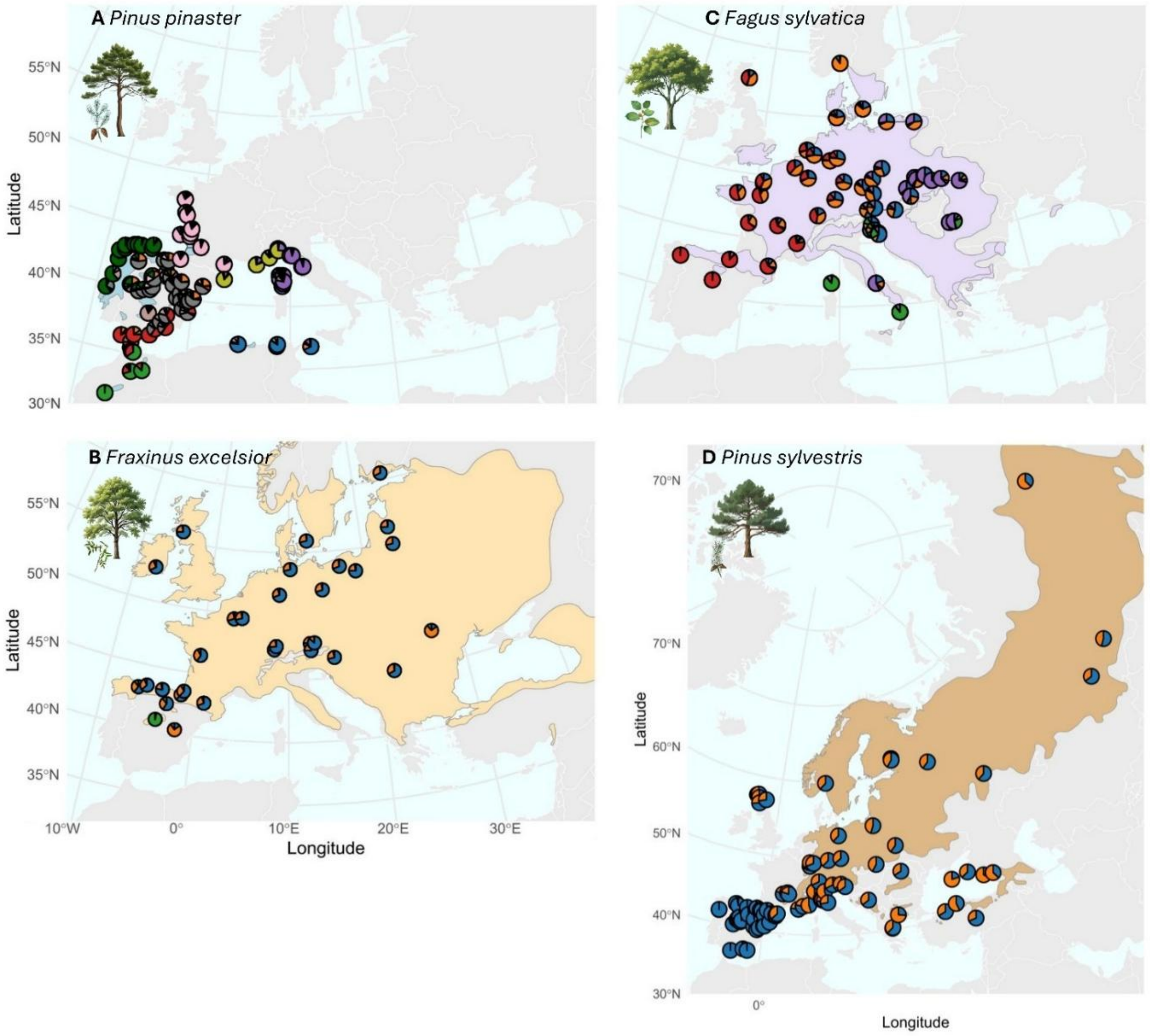


*Figure S7.* **Population genetic clusters inferred using STRUCTURE**. Panels A-D show the genetic clusters of *P. pinaster*, *F. excelsior*, *F. sylvatica*, and *P. sylvestris*, respectively. The genetic structure for *T. baccata* and *P. pinea* can be found in Francisco *et al*.[12] and Olsson *et al*.[14], respectively. Pie charts represent population-level ancestry proportions inferred from STRUCTURE analyses, with colours indicating the relative contribution of each genetic cluster. Shaded areas depict the approximate species ranges extracted Caudullo *et al*.[13], corresponding to the geographic extent of latitude 30°N-60°N and longitude 10°W-37°E, except for *P. sylvestris*, for which several sampled populations fall outside this range. The inferred number of genetic clusters varied markedly across species: two for *T. baccata* and *P. sylvestris*, three for *F. excelsior*, five for *F. sylvatica*, ten for *P. pinaster*, and twelve for *P. pinea*.

### *References*

14. Olsson, S., Jaramillo-Correa, J. P., Vajana, E., Pinosio, S., Macaya-Sanz, D., Bagnoli, F., Avanzi, C., Piotti, A., Alía, R., Mutke, S., Vendramin, G. G., Grivet, D., González-Martínez, S. C. Thriving with low diversity: demographic history and evolutionary dynamics of stone pine (*P. pinea* L). In preparation.
